# Nitrogen sharing strategies in six clonal species

**DOI:** 10.1101/2024.07.12.603230

**Authors:** Jana Duchoslavová

## Abstract

Nitrogen is often a limiting factor for plant growth, and its availability is a major determinant of level of competition. In clonal plants, patterns of nitrogen translocation between ramets may be part of plant nitrogen economics, and, as such, may also be related to the typical availability of nitrogen. In nutrient-poor habitats, extensive nutrient sharing balancing resource availability may be important, whereas nutrient sharing between established ramets may not be beneficial in productive habitats.

I tested the proposed nutrient sharing strategies on nitrogen translocation in six stoloniferous species that occur in habitats of varying productivity. Mother and daughter ramets of each species were grown either in a homogeneous nutrient-poor treatment or in a “nutrient-poor to nutrient-rich” treatment. I traced the translocation of nitrogen in both directions using stable isotope labelling when the daughter ramets were one month old.

Surprisingly, I found no effect of nutrient treatment on nitrogen translocation. Instead, each species translocated nitrogen either acropetally, basipetally, or equally in both directions. There was no relationship between the direction of translocation and the productivity of the species’ habitats. However, net translocation seemed to be related to the relative size of daughters across species, and within *Veronica officinalis*.

The results suggest that the relative size of plant parts is an important determinant of the strength of the sink for nitrogen they form, and that the growth habit of a species can affect its nitrogen translocation. Under certain conditions, such internally induced source-sink relationships may dominate over external nitrogen heterogeneity. I speculate that growth habit, together with nitrogen translocation patterns, may be part of adaptive growth strategies.

## Introduction

As autotrophic organisms, plants obtain both carbon and mineral nutrients for their growth directly from the abiotic environment. Mineral nutrients are taken up from the soil by plant roots and distributed within the plant to sites of use or storage (Tegeder and Masclaux-Daubresse 2018). The distribution of mineral nutrients in soils is highly heterogeneous even at fine spatial scales (Jackson and Caldwell 1993; Farley and Fitter 1999; Březina *et al.* 2019; Skálová *et al.* 2023) and plants explore and exploit this heterogeneity through their root systems (Giehl and von Wirén 2014; Weiser *et al.* 2016). In addition, environmental nutrient heterogeneity can be explored through clonal growth, which adds another level of plant modularity. Clonal plants, which grow laterally and form multiple rooting points along their horizontal stems, are able to acquire mineral nutrients at different sites and translocate them throughout the plant body (Noble and Marshall 1983; Alpert 1991; de Kroon *et al.* 1998). In contrast to the transport of nutrients between roots and shoots, this translocation is not inevitable because the rooting units (hereafter ramets) are potentially independent. Consequently, the extent of such clonal integration varies both between and within species (Alpert 1999; Si *et al.* 2020; Zhang *et al.* 2022).

Nitrogen is the most abundant mineral nutrient in plant tissues and is essential for many metabolic processes, including photosynthesis and nutrient uptake. In a non-clonal plant or a ramet of a clonal plant, nitrogen uptake, assimilation and distribution are regulated in a complex way and generally determined by source-sink relationships (Tegeder and Masclaux-Daubresse 2018). In a clonal plant with multiple ramets, such regulation and nitrogen distribution likely depend on level of ramet integration. In a highly integrated clonal plant, the nitrogen translocation may also be primarily determined by source-sink relationships affected by environmental heterogeneity (Evans 1991) and ramet size (Dong *et al.* 2015). Alternatively, nitrogen translocation may be unidirectional from older to younger parts (i.e. acropetal; Slade and Hutchings, 1987), possibly due to hormonal control (Alpert *et al.* 2002), or it may be generally limited. Across species or genotypes, translocation from younger to older parts (i.e. basipetal) seems to be the most variable (Evans, 1991; Lotscher and Hay, 1997; Slade and Hutchings, 1987, Duchoslavová and Jansa, unpubl.).

In natural environments, nitrogen is often a limiting factor for plant growth, and its availability is a major determinant of community composition and level of competition (Baer *et al.* 2004; Clark *et al.* 2007; Gough *et al.* 2012). Plants adapt to experienced nitrogen availability by adjusting the economy of its uptake, processing and conservation (Vázquez De Aldana and Berendse 1997). It is therefore an important component of a plant economics spectrum ranging from species adapted to low resource levels, focusing on resource conservation, to species of highly productive habitats, focusing on rapid resource acquisition and competition (Reich 2014). In clonal plants, patterns of nitrogen translocation between ramets may be part of plant nitrogen economics, and, as such, may also be related to the typical availability of nitrogen. Accordingly, the benefits of resource sharing were predicted to depend on environmental conditions (Hutchings and Price 1993; Gardner and Mangel 1999; Mágori and Oborny 2003).

In nutrient-poor habitats, extensive nutrient sharing to balance resource availability may be particularly important to maintain established ramets and to capture soil resources from a larger area, analogous to the ‘conservation’ end of the plant economics spectrum. Therefore, basipetal nutrient translocation may be beneficial in such environments where resource availability is generally low and unpredictable (Evans 1991). On the other hand, nutrient sharing between established ramets may not be beneficial in productive habitats where mineral nutrients are not limiting and competition for light is the main determinant of plant growth. Under such conditions, local nutrient availability may be used for rapid local growth and little nutrient sharing between established ramets may occur, analogous to the ‘acquisition’ end of the plant economics spectrum.

I aimed to test the proposed nutrient sharing strategies on translocation of nitrogen in six species that form aboveground horizontal stems and occur in habitats of varying productivity. Growth experiments examining the effect of translocation between ramets usually integrate the effect over a longer growth period and therefore do not separate the effect of early acropetal support and translocation between established ramets. I therefore chose the stable isotope labelling approach, which allows translocation to be examined in a short period of time during at a later stage of ramet development. Specifically, I hypothesised that (i) there would be no directional nitrogen translocation under homogeneous conditions, and (ii) species from nutrient-poor habitats would translocate nitrogen basipetally to ramets with lower nutrient availability, whereas species from productive habitats would not translocate nitrogen between established ramets.

## Methods

I used six stoloniferous species of non-forest habitats from different families, covering a gradient of habitat productivity measured by mean height of surrounding vegetation in database plots and Ellenberg’s N (Table 1; Chytrý et al., 2018; Herben et al., 2016). I grew each plant in a pair of 2 L pots, with the older part of the clone in one pot and the younger part in the other. In some species, these parts corresponded to clearly defined rooting ramets (*Hieracium bauhini, Fragaria viridis, Ranunculus repens, Potentilla reptans*), in others such a distinction was not possible due to the growth pattern of the species forming creeping monopodial stems with axillary inflorescences (*Veronica officinalis, Trifolium repens*). In all cases, the older and younger parts are hereafter referred to as mothers and daughters.

**Table 1.**
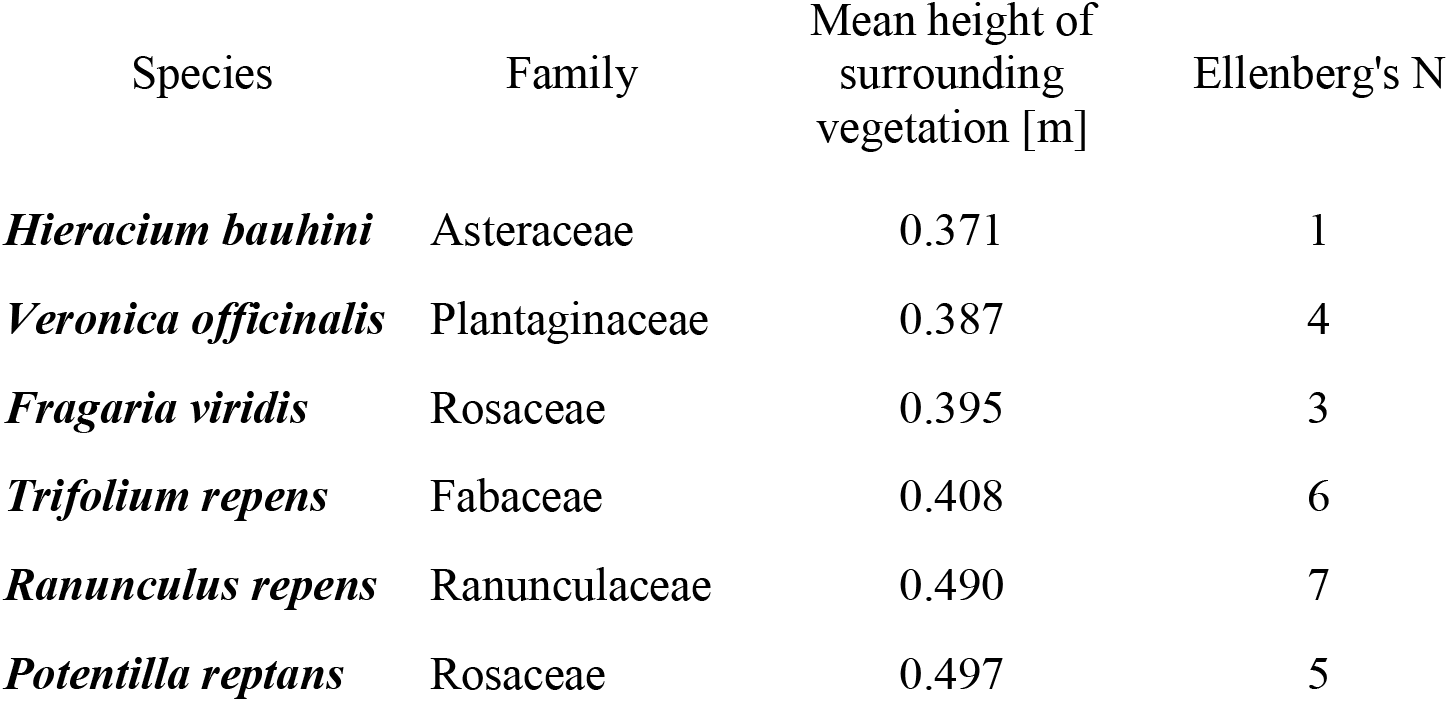
The species used in the experiment ordered according to mean height of surrounding vegetation in database plots.

Plant material originated from four genotypes per species, collected in the field (Central Bohemia, the Czech Republic). The mothers were potted in mid-June 2018 and dead plants were replaced till mid-July. Mothers were watered by solution of tap water and liquid NPK fertilizer (0.01 % Wuxal with 0.008 g N/l) in this preparation period. Rooting of daughters was initiated approximately 6 weeks after the planting of the mothers. The experimental nutrient regime was set at the time of initiation of daughter rooting. I used two nutrient treatments - ‘homogeneously poor’ and ‘poor to rich’ (0.025 % Wuxal with 0.02 g N/l for poor and 0.1 % Wuxal with 0.08 g N/l for rich conditions, Fig. 1).

**Figure 1.**
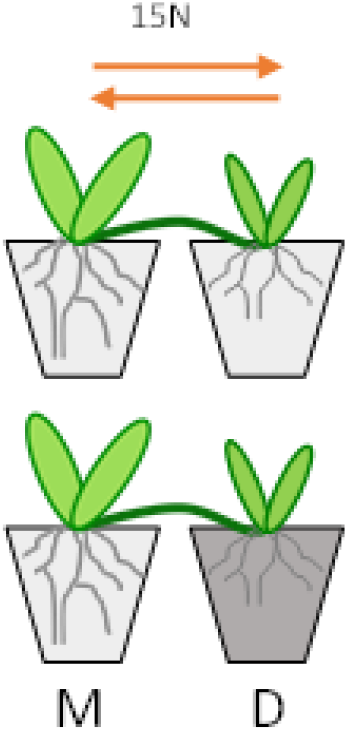
Experimental setup. Plants grew either in homogeneously poor or poor to rich conditions. Light colour of pot filling depicts low nutrient regime (0.02 % Wuxal with 0.016 g N/l), dark colour depicts rich nutrient regime (0.1 % Wuxal with 0.08 g N/l). The arrows indicate tracing of ^15^N translocation in both directions.

### Labelling and analyses

One month after daughter rooting initiation, I traced N translocation in both directions in each species and treatment, with four replicates for each combination of species, treatment, and direction. One extra plant per species was used for estimation of background ^15^N concentration.

^15^N was applied to the substrate of the pots with a syringe, at four positions per pot, 5 cm deep (total of 20 ml of ^15^NH_4_^15^NO_3_ solution per pot, 0.1 g/l). In half of the plants, the label was applied to the mother part and in the other half to the daughter part in order to trace the N translocation in both directions. The other part of the plants was treated with the same amount of unlabelled NH_4_NO_3_ solution. The labelling was conducted in four time blocks with one day difference, with all treatments represented in each block.

Two days after labelling, plants were harvested, connection between mother and daughter part was severed, shoots and roots were separated, and roots were carefully washed. After drying to constant weight (at 65°C), the dry weight of plant parts was estimated. The plant parts were then homogenised, ground to a fine powder in a ball mill (MM200, Retsch, Haan, Germany) and subjected to elemental and isotopic analysis. The N concentration and isotopic composition were measured using an elemental analyser (Flash EA 2000) coupled with an isotope ratio mass spectrometer (Delta V Advantage, ThermoFisher Scientific, Waltham, MA, USA).

I used following calculations to estimate the amount of ^15^N originating from pulse-labelling (i.e., excess ^15^N). First, F-ratios were calculated as *R*_*S*_/(*R*_*S*_ +1), where *R*_*S*_ stands for molar isotope ratio in a sample (^15^N/^14^N). The amount of total nitrogen (*C*, in moles) was then calculated as

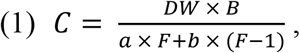

where *DW* is dry weight of a sample, *B* is molar concentration of nitrogen in a sample, *a* is equal to 14, *b* is equal to 15, and *F* stands for the F-ratio in a sample. The amount of ^15^N originating from pulse-labelling (*E*, in moles) was finally calculated as

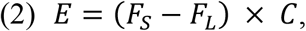

where F_S_ is F-ratio of a sample F_L_ is a limit F-ratio below which I cannot detect the stable-isotope enrichment with a sufficient confidence. This value was calculated from the F-ratios of the unlabelled control samples as 99^th^ percentile of a normal distribution. Therefore, F-ratio values above this limit would come from this distribution with probability less than 1 %.

The amount of ^15^N originating from pulse-labelling in unlabelled plant parts (roots and shoots combined) is further referred to as the amount of translocated ^15^N.

### Data analyses

I performed all the statistical analyses in the R statistical environment using linear models (R Core Team 2023). The biomass, root-to-shoot ratio and N uptake were log transformed in order to meet the model assumptions. Nitrogen translocation was analysed using a general model for all the species and additional separate models for each species to test for differences between the two directions of translocation.

Relationship of net nitrogen translocation and habitat productivity (measured as mean height of surrounding vegetation in database plots) and relationship of net nitrogen translocation and mean relative daughter size were modelled by simple linear regressions. In addition, separate models were performed to test for the effect of relative daughter size on nitrogen translocation to daughters within each species. Nitrogen translocation was again square-root transformed to meet the model assumptions.

## Results

### Comparison of species growth

There were marked differences among the sizes of mothers of different species, while the sizes of daughters, although significant, did not differ to such an extent (Fig. 2a, Table 2). Consequently, the species markedly differed in relative daughter size (i.e. daughter biomass to total biomass ratio, Fig. 2b).

**Figure 2.**
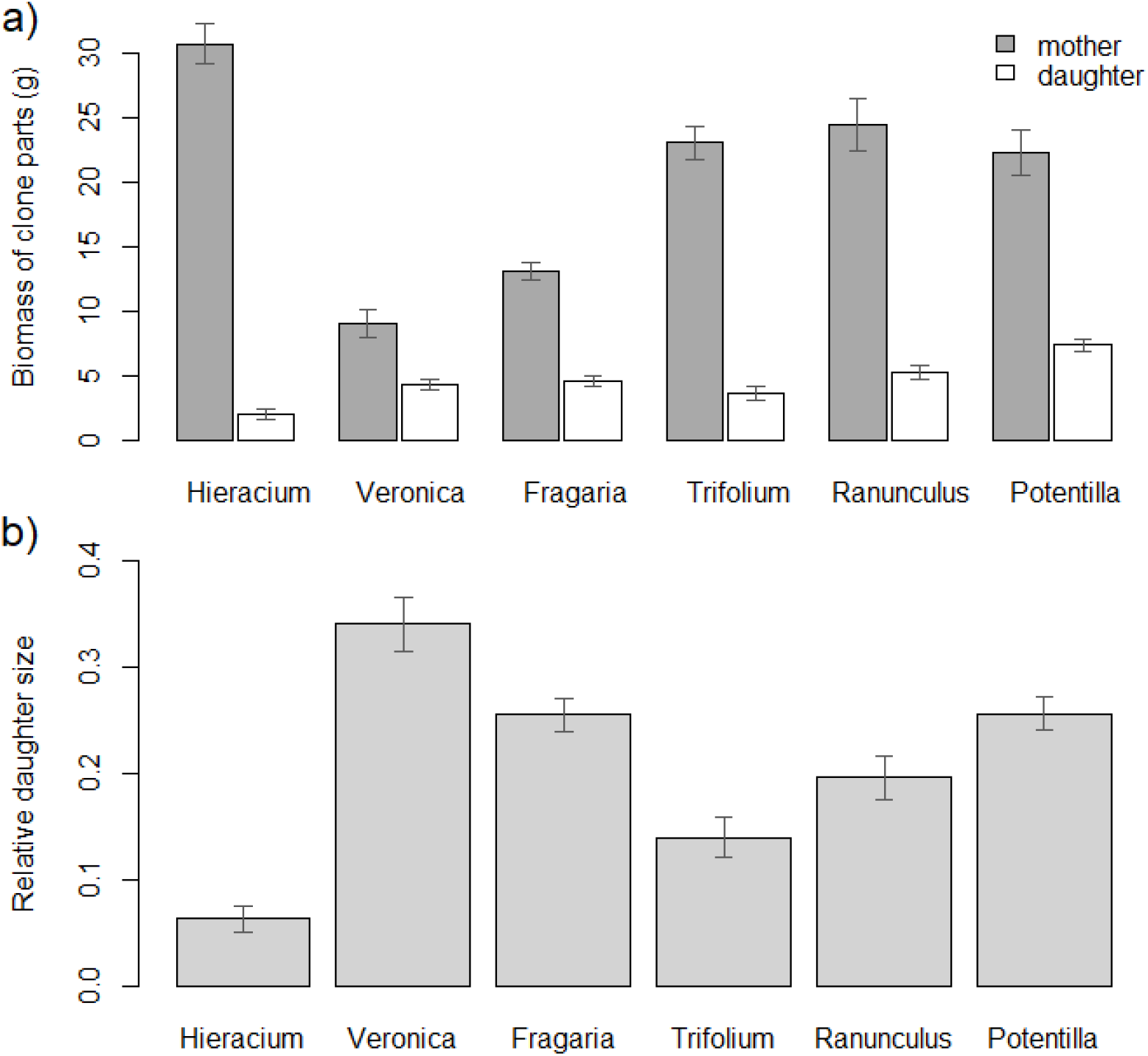
a) Biomass of mothers (grey bars) and daughters (white bars) and b) relative daughter size of the six species. Relative daughter size was calculated as daughter to total biomass ratio. Means are summarized across nutrient treatments; SEMs are depicted by arrows.

**Table 2.** ANOVA of linear models of biomass (log-transformed) of different species under the two nutrient treatments. Effects with P < 0.05 are highlighted in bold.

**Ramet biomass**
|  | d.f. | Biomass of daughters (log) |  |  | Biomass of mothers (log) |  |  |
| --- | --- | --- | --- | --- | --- | --- | --- |
|  |  | Sum Sq | *F* | *P* | Sum Sq | *F* | *P* |
| Species | **5** | **16.64** | **18.06** | **<0.001** | **18.70** | **33.21** | **<0.001** |
| Nutrients | **1** | **1.45** | **7.88** | **0.006** | 0.01 | 0.13 | 0.718 |
| Species x nutrients | 5 | 1.86 | 2.02 | 0.085 | 0.28 | 0.49 | 0.780 |
| Residuals | 82 | 15.12 |  |  | 9.23 |  |  |

Nutrient regime had a significant positive effect on biomass of daughters (*P*=0.006), with only marginally significant differences among species (*P*=0.085, Fig. 3, Table 2). There was no significant effect of the nutrient regime on mother biomass (*P*=0.718, Table 2).

**Figure 3.**
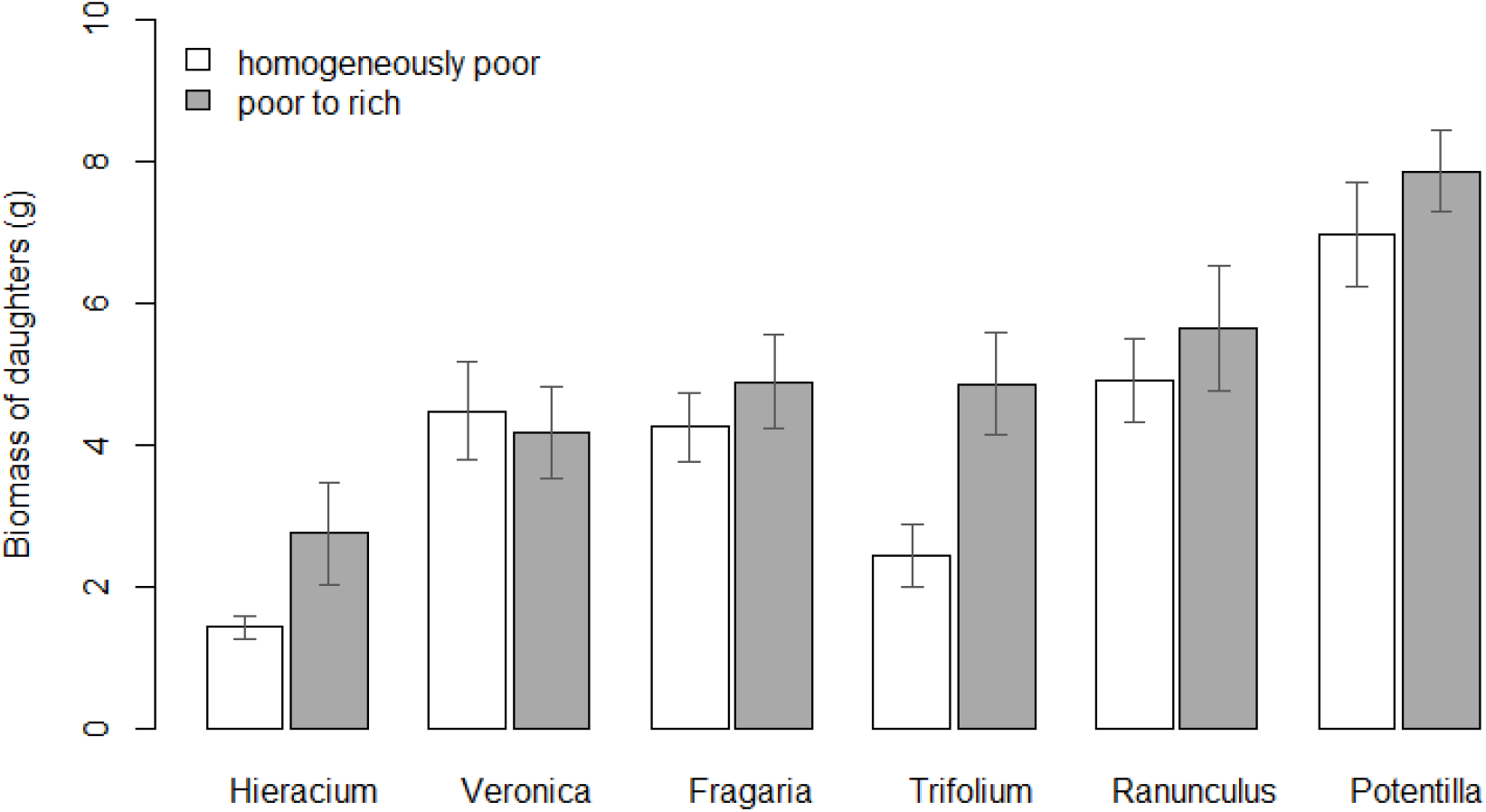
Effect of nutrient treatment on biomass of daughters. Means and SEMs depicted.

Root-to-shoot ratio significantly differed between mothers and daughters and the difference was species-specific (Table 3, Fig. 4). Whereas the root-to-shoot ratio of mothers and daughters was comparable in Fragaria and Trifolium, daughters had a markedly higher root-to-shoot ratio than mothers in Hieracium, and daughters had a lower root-to-shoot ratio than mothers in Veronica, Ranunculus and Potentilla. There was also a weak species-specific effect of nutrients on root-to-shoot ratio. Whereas most species had a neutral or negative response of root-to-shoot ratio to added nutrients, Trifolium had a higher root-to-shoot ratio in the poor-to-rich nutrient treatment (Fig. 4).

**Table 3.** ANOVA of a linear model of root-to-shoot ratio (log-transformed) of different species under the two nutrient treatments. Effects with P < 0.05 are highlighted in bold.

Root-shoot ratio of daughters and mothers (log)
|  | d.f. | Sum Sq | F | P |
| --- | --- | --- | --- | --- |
| **Species** | **5** | **8.78** | **11.50** | **<0.001** |
| **Mother/daughter** | **1** | **6.48** | **42.47** | **<0.001** |
| Nutrients | 1 | 0.00 | 0.01 | 0.908 |
| **Species x mother/daughter** | **5** | **13.92** | **18.24** | **<0.001** |
| **Species x nutrients** | **5** | **2.20** | **2.89** | **0.016** |
| Mother/daughter x nutrients | 1 | 0.10 | 0.64 | 0.426 |
| Species x mother/daughter x nutrients | 5 | 1.23 | 1.61 | 0.159 |
| Residuals | 164 | 25.03 |  |  |

**Figure 4.**
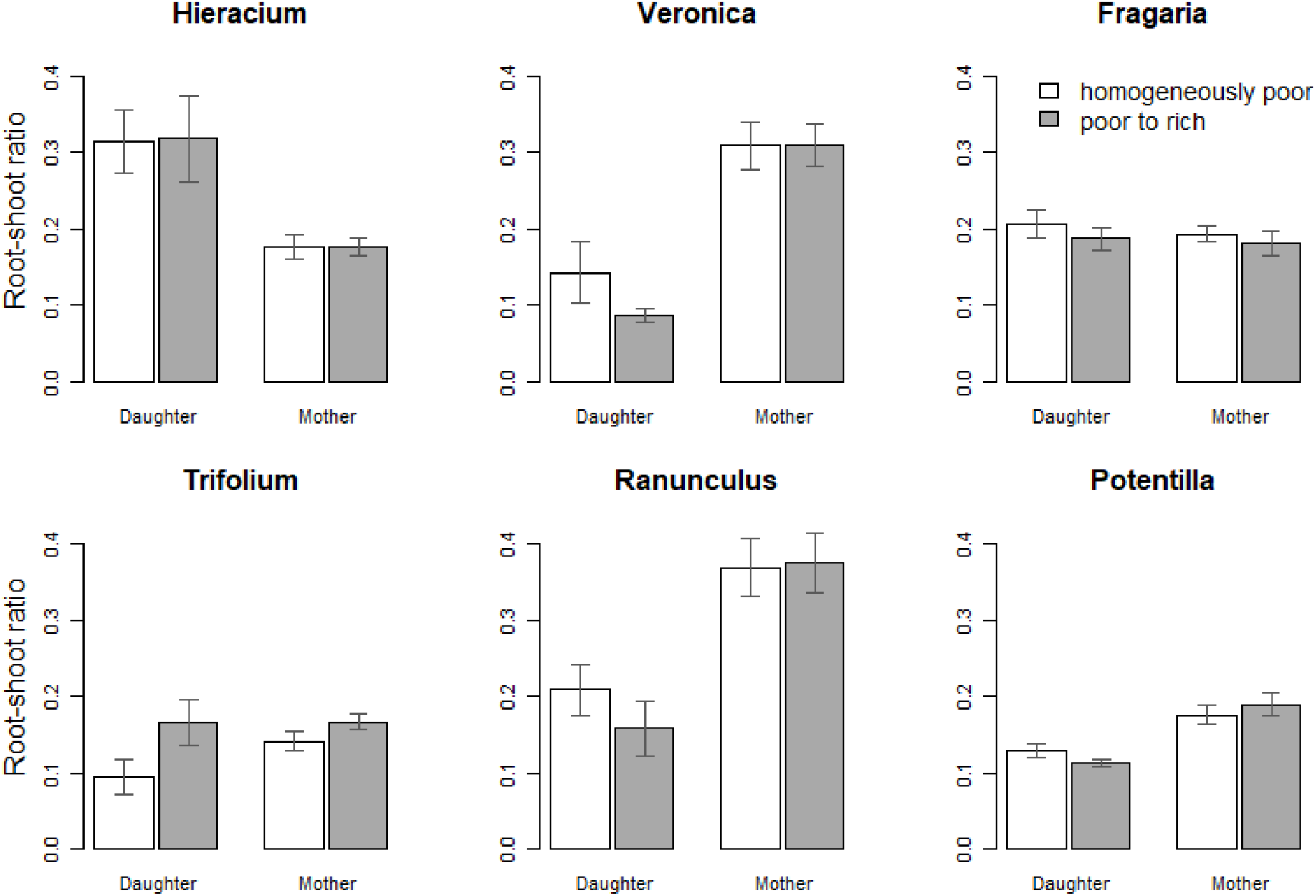
Root-to-shoot ratio of daughter and mother parts in homogeneously poor (white) and poor to rich treatment (grey).

### Nitrogen uptake

There were no marked differences in mothers’ nitrogen uptake between species (Fig. 5 – y axis) or treatments. N uptake of daughters was generally lower than the uptake of mothers and it differed significantly between species, with *Veronica* uptake markedly lower than uptake of all the other species (Table 4, Fig. 5 – x axis). This could be due to the very shallow root system of *Veronica* daughters, which may not reach the main volume of applied label. Therefore, the translocation of nitrogen to the mother may be somewhat underestimated in this species. Nitrogen treatment had no significant effect on nitrogen uptake.

**Table 4.** ANOVA of a linear model of nitrogen uptake (log-transformed) of different species and mother or daughter ramets under the two nutrient treatments. Effects with P < 0.05 are highlighted in bold.

| Nitrogen uptake of ramets (log) |  |  |  |  |
| --- | --- | --- | --- | --- |
|  | d.f. | Sum Sq | *F* | *P* |
| **Species** | **5** | **8.79** | **6.80** | **<0.001** |
| **Mother/daughter** | **1** | **13.08** | **50.58** | **<0.001** |
| Nutrients | 1 | 0.39 | 1.49 | 0.226 |
| **Species x mother/daughter** | **5** | **13.84** | **10.71** | **<0.001** |
| Species x nutrients | 5 | 1.07 | 0.83 | 0.533 |
| Mother/daughter x nutrients | 1 | 0.74 | 2.87 | 0.095 |
| Species x mother/daughter x nutrients | 5 | 0.97 | 0.75 | 0.587 |
| Residuals | 69 | 17.8433 |  |  |

**Figure 5.**
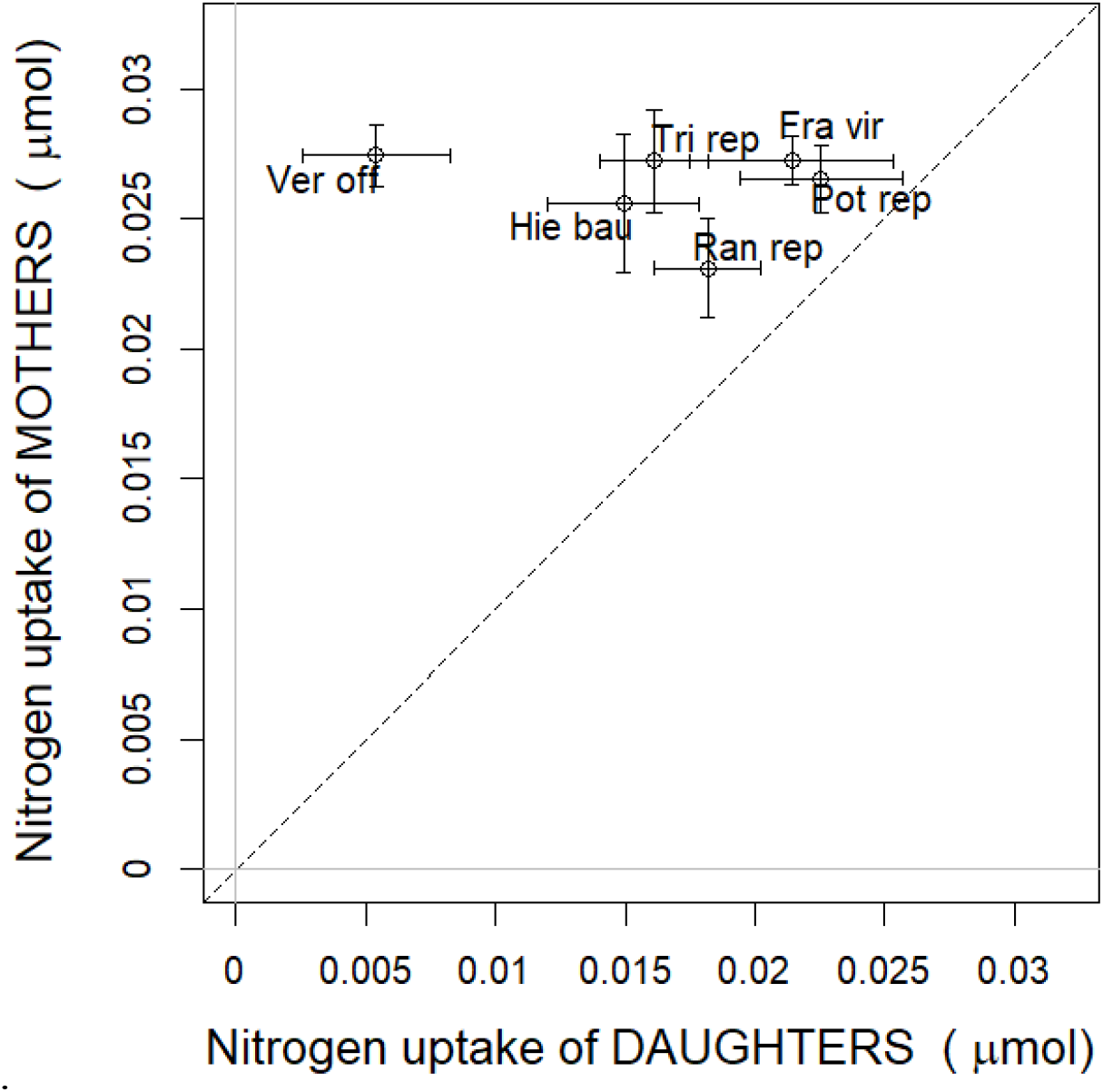
Nitrogen uptake of mothers (y axis) and daughters (x axis) of the six species. Dashed 1:1 line connects positions with equal uptake of daughters and mothers. Means and SEMs across nutrient treatments depicted.

### Nitrogen translocation

Direction of nitrogen translocation varied significantly between the species, with no significant effect of nutrient treatment (Table 5). Whereas *Veronica, Fragaria* and *Potentilla* translocated more nitrogen to daughters, *Ranunculus* and *Hieracium* translocated more nitrogen to mothers. The translocation of *Trifolium* did not differ significantly between the two directions, so net translocation was close to zero in this species (Fig. 6, Table 6).

**Table 5.** Translocated nitrogen (square-root transformed) of different species under the two nutrient treatments - ANOVA of a linear model. Direction of traced translocation reflects labelling of daughters or mothers. Effects with P < 0.05 are highlighted in bold.

|  | d.f. | Sum Sq | *F* | *P* |
| --- | --- | --- | --- | --- |
| **Species** | **5** | **0.006** | **4.131** | **0.002** |
| **Direction** | **1** | **0.003** | **8.721** | **0.004** |
| Nutrients | 1 | 0.000 | 0.160 | 0.690 |
| **Species x direction** | **5** | **0.033** | **21.107** | **<0.001** |
| Species x nutrients | 5 | 0.002 | 1.132 | 0.352 |
| Direction x nutrients | 1 | 0.000 | 1.421 | 0.237 |
| Species x direction x nutrients | 5 | 0.001 | 0.815 | 0.543 |
| Residuals | 69 | 0.021 |  |  |

**Table 6.**
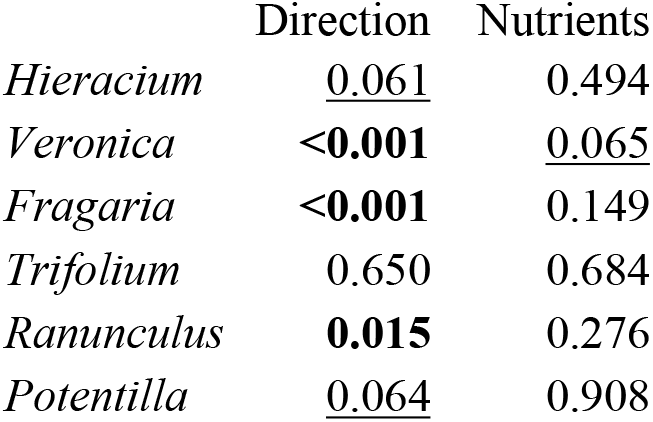
P-values for effects of direction and nutrients on translocation from linear models run separately for each species. Direction of traced translocation reflects labelling of daughters or mothers. Effects with P < 0.05 are highlighted in bold, marginally significant effects with P < 0.1 are underlined.

**Figure 6.**
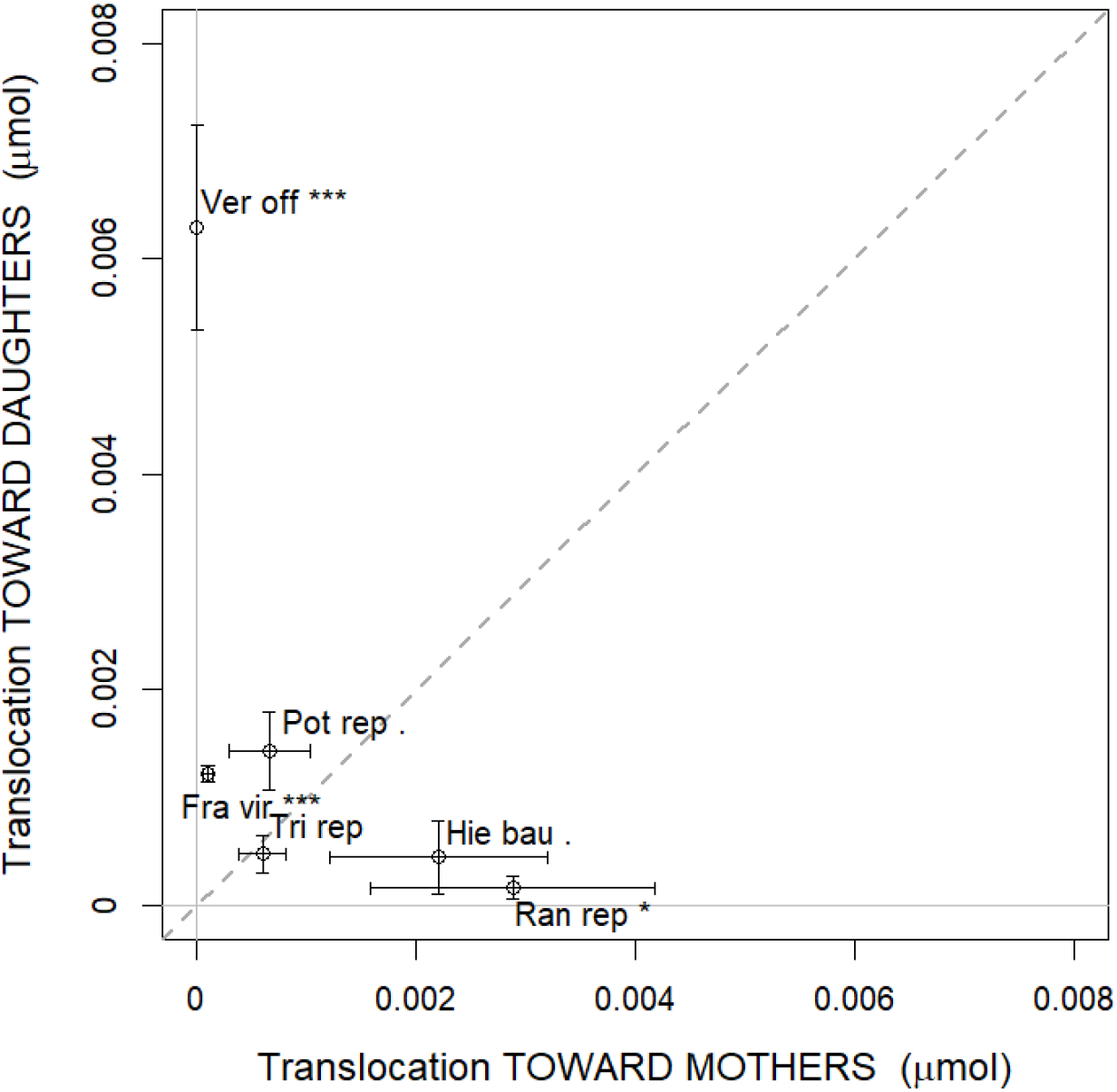
Two-way translocation plot (means and SEMs). Dashed 1:1 line connects positions with equal translocation in both directions (zero net translocation). Means and SEMs across nutrient treatments depicted.

*Veronica* was the only species that tended to respond to the nutrient treatment, with increased nitrogen translocation to daughters in high nutrient level (Table 6, Fig. 7).

**Figure 7.**
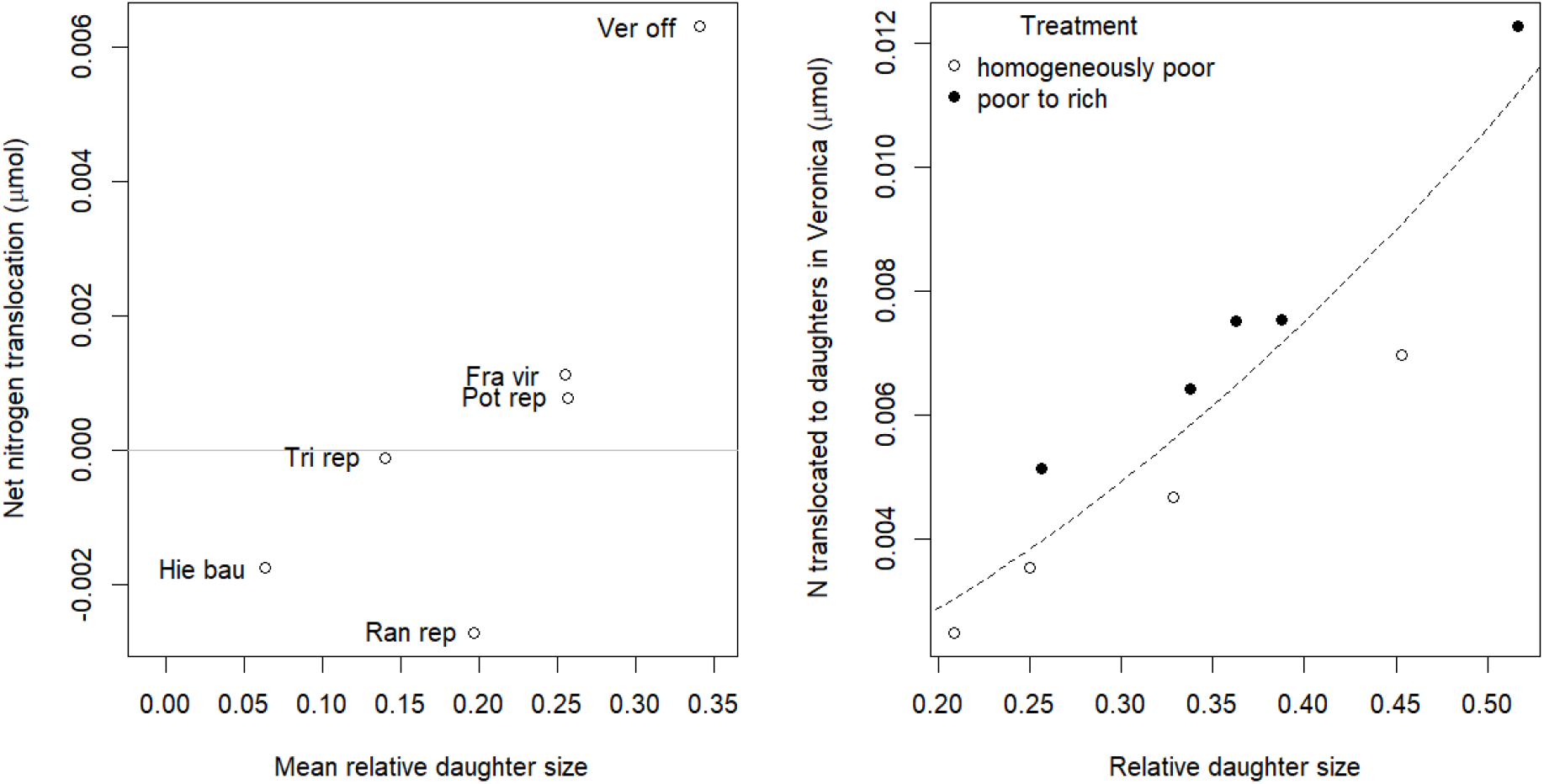
Relationship of net nitrogen translocation to mean relative daughter size (left) and relationship of nitrogen translocation to daughters to relative daughter size and nutrient treatment in Veronica (right).

There was no relationship between net nitrogen translocation and habitat productivity of the species (*P*=0.506, R^2^=0.12, n=6).

### Nitrogen translocation and relative daughter size

Net nitrogen translocation tended to increase with mean relative daughter size of the species (*P*=0.063, R^2^=0.62, n=6; Fig. 7). Furthermore, there was a relationship between the nitrogen translocated to daughters and relative daughter size in *Veronica* (*P* < 0.001, R^2^=0.83; Fig. 7). No such relationship was observed in the other species.

## Discussion

By tracing of ^15^N in both directions, I found distinct patterns of nitrogen translocation between mothers and established daughters in the six stoloniferous species. Surprisingly, the translocation was not affected by the nutrient regime of the daughters. Irrespective of nutrient treatment, three species translocated nitrogen predominantly to daughter parts, two species to mother parts and one species showed zero net translocation of nitrogen between mothers and daughters. To our knowledge, this is the first attempt to examine translocation strategies using a comparative approach (but see e.g. Ashmun et al., 1982 or Xu et al., 2010 for two species comparisons).

I hypothesised that nitrogen would not be translocated under the homogeneous conditions and that the nitrogen sharing strategy under heterogeneous nutrient availability would reflect the productivity of the habitats typically experienced by the species. Our results did not support these hypotheses. It is possible that the number of species and the length of the productivity gradient used in our study were not sufficient to show the pattern. In general, there is a lot of variability within communities and functional groups, although there are visible patterns of plant economic strategies along environmental gradients (Maire et al., 2009).

Regardless of the environmental heterogeneity, the species studied differed greatly in their growth habit. These differences resulted in different allocations to mother and daughter parts, affecting the relative size of mother and daughter roots (i.e. sources of nutrients) and their shoots, which presumably determine the strength of nutrient sinks.

The relative allocation to acquisition of soil resources is indicated by the root-to-shoot ratio, which, in daughter ramets of stoloniferous species, inevitably increases with the development. Accordingly, the acropetal nutrient translocation is highly important for the growth of new ramets that are developing their own roots (Marshall 1990; Dong *et al.* 2015). However, the root-to-shoot ratio may remain lower in daughters than in mothers in some species due to a “developmentally programmed division of labour”, in which daughters specialise in the acquisition of photosynthates and mothers support them with nutrients (Roiloa 2019; Xi *et al.* 2019). Therefore, prevailing translocation to daughters could be expected especially if the daughter root-to-shoot ratio is low. At the time of harvest, root-to-shoot ratio was lower in daughters than in mothers of four species, comparable between daughters and mothers in *Fragaria* and higher in daughters in *Hieracium*. However, there was no clear link between the relative root-to-shoot ratio values and net nitrogen translocation in the species.

The relative size of daughters may reflect the strength of the sink for the nutrients that they form. The results indeed indicated that net translocation of nitrogen was directed to the relatively larger daughters. This relationship was driven particularly by two of the species – by *Hieracium* forming relatively small daughters which translocate nitrogen basipetally to mothers, and by *Veronica* with relatively large daughters and high acropetal nitrogen translocation. Moreover, I observed the relationship between daughter relative size and the magnitude of translocation to daughters also at the intraspecific level in *Veronica*. Surprisingly, higher nutrient availability to daughters seemed to increase nitrogen translocation to them in this species. This pattern resembles the ‘rich get richer’ effect proposed in *Fragaria chiloensis* (Alpert 1996) and *Buchloe dactyloides* (Sun *et al.* 2011).

Our results suggested that the growth habit of clonal species, and in particular the relative size of their ramets, is more likely to determine nutrient translocation than environmental heterogeneity in nutrient availability. The observed distinct patterns of nutrient translocation may be either an unavoidable consequence of biomass allocation or, together with biomass allocation, part of adaptive resource-sharing strategies. I hypothesise that the acropetal nitrogen translocation may facilitate exploration of new patches, whereas daughters may serve as extended hands for nitrogen acquisition in plants with the prevailing basipetal translocation (Pinno and Wilson 2014; Duchoslavová and Jansa 2018). Such function of daughter ramets may be temporal and serve, for example, to support flowering mother ramets by vegetative daughters, as could be the case of *Hieracium* in our experiment. In contrast to the nitrogen translocation observed here, carbon translocation seems to respond more readily to gradients in light availability and different carbon translocation strategies are not pronounced under homogeneous conditions (Duchoslavová and Jansa, 2018; Duchoslavová and Jansa, unpubl.).

More information on the net translocation of nutrients between mother and daughter parts of different clonal plants is needed to generalise the results. As growth experiments do not separate the effect of early translocation from translocation between established ramets, I encourage future studies to examine translocation of nutrients in both directions by the labelling approach, which has rarely been done to date (Pinno and Wilson 2014; Duchoslavová and Jansa 2018; Dong *et al.* 2022).

## Acknowledgements

I thank to Tomáš Herben and Jitka Klimešová for their valuable comments, and Jan Jansa for his help with the isotopic and elemental analyses.

## References

Alpert P. 1991. Nitrogen sharing among ramets increases clonal growth in Fragaria chiloensis. Ecology 72: 69–80.

Alpert P. 1996. Nutrient sharing in natural clonal fragments of Fragaria chiloensis. Journal of Ecology 84: 395–406.

Alpert P. 1999. Clonal integration in Fragaria chiloensis differs between populations: ramets from grassland are selfish. Journal of Ecology 120: 69–76.

Alpert P, Holzapfel C, Benson J. 2002. Hormonal modification of resource sharing in the clonal plant Fragaria chiloensis. Functional Ecology 16: 191–197.

Ashmun JW, Thomas RJ, Pitelka LF. 1982. Translocation of photoassimilates between sister ramets in two rhizomatous forest herbs. Annals of Botany 49: 403–415.

Baer SG, Blair JM, Collins SL, Knapp AK. 2004. Plant community responses to resource availability and heterogeneity during restoration. Oecologia 139: 617–629.

Březina S, Jandová K, Pecháčková S, et al. 2019. Nutrient patches are transient and unpredictable in an unproductive mountain grassland. Plant Ecology 220: 111–123.

Chytrý M, Tichý L, Dřevojan P, Sádlo J, Zelený D. 2018. Ellenberg-type indicator values for the Czech flora. Preslia 90: 83–103.

Clark CM, Cleland EE, Collins SL, et al. 2007. Environmental and plant community determinants of species loss following nitrogen enrichment. Ecology Letters 10: 596–607.

Dong B, Alpert P, Zhang Q, Yu F. 2015. Clonal integration in homogeneous environments increases performance of Alternanthera philoxeroides. Oecologia 179: 393–403.

Dong BC, Wang P, Luo FL. 2022. Sharing of nitrogen between connected ramets of Alternanthera philoxeroides in homogeneous environments. Plant and Soil 478: 445–460.

Duchoslavová J, Jansa J. 2018. The direction of carbon and nitrogen fluxes between ramets in Agrostis stolonifera changes during ontogeny under simulated competition for light. Journal of Experimental Botany 69: 2149–2158.

Evans JP. 1991. The effect of resource integration on fitness related traits in a clonal dune perennial, Hydrocotyle bonariensis. Oecologia 86: 268–275.

Farley RA, Fitter AH. 1999. Temporal and spatial variation in soil resources in a deciduous woodland. Journal of Ecology 87: 688–696.

Gardner SN, Mangel M. 1999. Modeling investments in seeds, clonal offspring, and translocation in a clonal plant. Ecology 80: 1202–1220.

Giehl RFH, von Wirén N. 2014. Root nutrient foraging. Plant Physiology 166: 509–517.

Gough L, Gross KL, Cleland EE, et al. 2012. Incorporating clonal growth form clarifies the role of plant height in response to nitrogen addition. Oecologia 169: 1053–62.

Herben T, Chytrý M, Klimešová J. 2016. A quest for species-level indicator values for disturbance. Journal of Vegetation Science 27: 628–636.

Hutchings MJ, Price EAC. 1993. Does Physiological Integration Enable Clonal Herbs to Integrate the Effects of Environmental Heterogeneity? Plant Species Biology 8: 95–105.

Jackson R, Caldwell M. 1993. The scale of nutrient heterogeneity around individual plants and its quantification with geostatistics. Ecology 74: 612–614.

de Kroon H, van der Zalm E, van Rheenen J, van Dijk A, Kreulen R. 1998. The interaction between water and nitrogen translocation in a rhizomatous sedge (Carex flacca). Oecologia 116: 38–49.

Lotscher M, Hay MJM. 1997. Genotypic differences in physiological integration, morphological plasticity and utilization of phosphorus induced by variation in phosphate supply in Trifolium repens. The Journal of Ecology 85: 341.

Mágori K, Oborny B. 2003. Cooperation and competition in heterogeneous environments: the evolution of resource sharing in clonal plants. Evolutionary Ecology Research 5: 787–817.

Marshall C. 1990. Source-sink relations of interconnected ramets In: van Groenendael J, de Kroon H, eds. Clonal growth in plants: regulation and function. The Hague: SPB Academic Publishing, 23–41.

Noble J, Marshall C. 1983. The population biology of plants with clonal growth: II. The nutrient strategy and modular physiology of Carex arenaria. The Journal of Ecology 71: 865– 877.

Pinno BD, Wilson SD. 2014. Nitrogen translocation between clonal mother and daughter trees at a grassland–forest boundary. Plant Ecology 215: 347–354.

R Core Team. 2023. R: A language and environment for statistical computing.

Reich PB. 2014. The world-wide “fast-slow” plant economics spectrum: A traits manifesto. Journal of Ecology 102: 275–301.

Roiloa SR. 2019. Clonal traits and plant invasiveness: The case of Carpobrotus N.E.Br. (Aizoaceae). Perspectives in Plant Ecology, Evolution and Systematics 40: 125479.

Si C, Alpert P, Zhang JF, et al. 2020. Capacity for clonal integration in introduced versus native clones of the invasive plant Hydrocotyle vulgaris. Science of the Total Environment 745: 141056.

Skálová H, Jandová K, Balšánková T, et al. 2023. Cations make a difference: Soil nutrient patches and fine-scale root abundance of individual species in a mountain grassland. Functional Ecology 37: 915–928.

Slade AJ, Hutchings MJ. 1987. An analysis of the costs and benefits of physiological integration between ramets in the clonal perennial herb Glechoma hederacea. Oecologia 73: 425–431.

Sun X-L, Niu J-Z, Zhou H. 2011. Buffalograss decreases ramet propagation in infertile patches to enhance interconnected ramet proliferation in fertile patches. Flora - Morphology, Distribution, Functional Ecology of Plants 206: 380–386.

Tegeder M, Masclaux-Daubresse C. 2018. Source and sink mechanisms of nitrogen transport and use. New Phytologist 217: 35–53.

Vázquez De Aldana BR, Berendse F. 1997. Nitrogen-use efficiency in six perennial grasses from contrasting habitats. Functional Ecology 11: 619–626.

Weiser M, Koubek T, Herben T. 2016. Root foraging performance and life-history traits. Frontiers in Plant Science 7: 1–10.

Xi DG, You WH, Hu AA, Huang P, Du DL. 2019. Developmentally programmed division of labor in the aquatic invader Alternanthera philoxeroides under homogeneous soil nutrients. Frontiers in Plant Science 10: 1–9.

Xu C, Schooler S, van Klinken R. 2010. Effects of clonal integration and light availability on the growth and physiology of two invasive herbs. Journal of Ecology 98: 833–844.

Zhang XM, He LX, Xiao X, et al. 2022. Clonal integration benefits an invader in heterogeneous environments with reciprocal patchiness of resources, but not its native congener. Frontiers in Plant Science 13: 1–10.

